# Surface temperatures are influenced by handling stress independently of corticosterone levels in wild king penguins *(Aptenodytes patagonicus*)

**DOI:** 10.1101/2023.06.16.545254

**Authors:** Agnès Lewden, Chelsea Ward, Aude Noiret, Sandra Avril, Lucie Abolivier, Caroline Gérard, Tracey L. Hammer, Émilie Raymond, Jean-Patrice Robin, Vincent A. Viblanc, Pierre Bize, Antoine Stier

## Abstract

Assessing the physiological stress responses of wild animals opens a window for understanding how organisms cope with environmental challenges. Since stress response is associated with changes in body temperature, the use of body surface temperature through thermal imaging could help to measure acute and chronic stress responses non-invasively. We used thermal imaging, acute handling-stress protocol and an experimental manipulation of corticosterone (the main glucocorticoid hormone in birds) levels in breeding king penguins (*Aptenodytes patagonicus*), to assess: 1. the potential contribution of the Hypothalamo-Pituitary-Adrenal (HPA) axis in mediating chronic and acute stress-induced changes in adult surface temperature, 2. the influence of HPA axis manipulation on parental investment through thermal imaging of eggs and brooded chicks, and 3. the impact of parental treatment on offspring thermal’s response to acute handling.

Maximum eye temperature (*T_eye_*) increased and minimum beak temperature (*T_beak_*) decreased in response to handling stress in adults, but neither basal nor stress-induced surface temperatures were significantly affected by corticosterone implant. While egg temperature was not significantly influenced by parental treatment, we found a surprising pattern for chicks: chicks brooded by the (non-implanted) partner of corticosterone-implanted individuals exhibited higher surface temperature (both *T_eye_* and *T_beak_*) than those brooded by glucocorticoid-implanted or control parents. Chick’s response to handling in terms of surface temperature was characterized by a drop in both *T_eye_* and *T_beak_* independently of parental treatment.

We conclude that the HPA axis seems unlikely to play a major role in determining chronic or acute changes in surface temperature in king penguins. Changes in surface temperature may primarily be mediated by the Sympathetic-Adrenal-Medullary (SAM) axis in response to stressful situations. Our experiment did not reveal a direct impact of parental HPA axis manipulation on parental investment (egg or chick temperature), but a potential influence on the partner’s brooding behaviour.

**Highlights:**

- Experimental increase in corticosterone does not affect king penguin’s surface temperature
- Acute handling stress increases eye but decreases beak surface temperature in adults
- Acute handling decreases both eye and beak surface temperatures in young chicks
- Parental corticosterone treatment does not affect egg surface temperature during incubation
- Chicks brooded by non-implanted partners of corticosterone parents are warmer than chicks brooded by others adults.

## 1. Introduction

Measuring stress response is of central interest in wild animals to understand how they cope with environmental change (Ellenberg et al., 2007; Romero, 2004). Traditionally, ecologists have evaluated acute stress by assessing the activation of the Sympathetic-Adrenal-Medullary (SAM) or Hypothalamic-Pituitary-Adrenal (HPA) axes, either at baseline levels or in response to acute disturbances in the environment, by metrics such as increased heart rate (Cabanac and Guillemette, 2001; Viblanc et al., 2012), ventilation rate (Carere and Oers, 2004) or glucocorticoid hormone (GC) secretion (Sapolsky et al., 2000). Stress exposure is also known to trigger core body and surface temperature changes (Oka, 2018). As the latter can be measured using thermal imaging (McCafferty, 2013), it led to a recent burgeoning of thermal imaging studies to measure the stress response of captive and wild endotherms (McCafferty et al., 2021). Both the SAM and HPA axes can potentially influence changes in body surface temperature, but the importance of these two pathways in mediating the stress response(s) measured through thermal imaging remains largely unknown (but see Jerem and Romero (2023)). Considering that both HPA and SAM axes provide strong and very reactive responses to stress, teasing apart their contribution to changes in peripheral temperature is challenging.

SAM axis responds within seconds to a stressor by releasing catecholamines hormones (adrenaline and noradrenaline) into the blood stream, which induces an immediate increase in heart rate (tachycardia) and vasoconstriction of peripheral blood vessels (Wingfield and Romero, 2015). This redistribution of peripheral blood flow to essential internal organs generally leads to a response called stress-induced hyperthermia along with a decrease of surface temperature (Cabanac and Guillemette, 2001; Herborn et al., 2015; Oka, 2018). The SAM axis response is followed by the slower activation of HPA axis that releases GC into the blood stream within a few minutes (Wingfield and Romero, 2015). GC are metabolic hormones that helps responding to a stressor by mobilizing energy resources and by triggering an increase in behavioural activity and metabolic rate (Wingfield and Romero, 2015). Circulating GC levels are usually considered at baseline (*i.e.* without an inducing stressor) or at stress-induced levels, which are often interpreted as mirroring chronic stress/metabolic demand and acute stress response respectively. Yet, the actual interpretation of circulating GC levels is likely more complex (MacDougall-Shackleton et al., 2019). GC-mediated changes in metabolism may cause an increase in internal heat production that, in turn, may lead to changes in body surface temperature (Oka, 2018). Additionally, previous evidence suggests that GC might be necessary for enabling the SAM axis to induce shivering, free fatty acid mobilization or vasoconstriction (Deavers and Musacchia, 1979).

Therefore, both the SAM and HPA axes could trigger, independently from each other, or together, a change in body core and/or surface temperatures (Ouyang et al., 2021). The immediate activation of SAM axis could lead to a decrease of peripheral temperature independently of GC release by the HPA axis. On the other hand, the short to long-term elevation of baseline GC might lead to increased metabolic rate, heat production and, ultimately, higher peripheral temperature (to facilitate heat dissipation and maintain homeothermy) independently of the SAM axis. The use of thermal imaging as a non-invasive tool to measure stress in unmanipulated animals is actively developing, but the implication of the SAM and HPA axes in mediating changes in surface temperature remains uncertain (Jerem and Romero, 2023; Ouyang et al., 2021). For instance, data in captive and wild birds show that acute stress exposure often leads to a decrease in body surface temperature (reviewed in Table 1, but note some discrepancies between species and/or body parts), as predicted if these changes are driven by the SAM axis. However, evidence for an impact of chronic stressors on body surface temperature (Herborn et al., 2018; Robertson et al., 2020a, 2020b; Winder et al., 2020) or for a correlation between circulating GC levels and body surface temperature (Giloh et al., 2012; Jerem et al., 2018; Weimer et al., 2020) remains unclear.

**Table 1:** Summary table of avian studies testing the impact of acute stress exposure on changes in body surface temperatures measured with thermal imaging. (studies related to heat and food stress are not included). *Area* corresponds to the body region being measured, response to the direction of the temperature response (=: no significant change, ↓: significant decrease and ↑: significant increase, *Δ T°C* to the amplitude of the temperature response, *stressor* the nature and duration of the experimental stressor. Some studies measured temperature responses at different time points, and when the effects of stress exposure change through time, the responses are shown with consecutive symbols (*e.g.* =, ↑: initial absence of change followed later on by an increase in surface temperature). Please note that the area of interest for T_eye_ differs between studies (periorbital region *vs.* eyeball region) and that the surface temperature being measured can differ in nature between studies (*i.e.* maximum, average or minimum temperature of the area of interest).

| Species | Condition | Age | Stressor | Area | Response | $\Delta T^{\circ}\text{C}$ | Ref. |
| --- | --- | --- | --- | --- | --- | --- | --- |
| Blue tit<br>*Cyaniste caeruleascens* | Wild | Adult | Trap closure +<br>handling<br>(3min) | Eye | ↓, ↑, = | -1.0, +1.0°C, = | (Jerem et al., 2019) |
| Budgerigars<br>*Melopsittacus undulatus* | Captive | Adult | Handling<br>(30 min) | Eye<br>Legs | ↑, =<br>= | +0.7°C, =<br>= | (Ikkatai and<br>Watanabe, 2015) |
| Chicken<br>*Gallus gallus domesticus* | Captive | Adult | Air puff<br>(10 min) | Eye | ↓ | -1.0°C | (Edgar et al., 2011) |
| Chicken<br>*Gallus gallus domesticus* | Captive | Adult | Handling<br>(20 min) | Eye<br>Head<br>Comb | ↓, =<br>=, ↑<br>↓, = | -0.5°C, =<br>=, +1.0°C<br>-1.5°C, = | (Edgar et al., 2013) |
| Chicken<br>*Gallus gallus domesticus* | Captive | Chick<br>(9 weeks) | Air puff<br>(10 min) | Eye | ↓ | -1.0°C | (Edgar and Nicol,<br>2018) |
| Chicken<br>*Gallus gallus domesticus* | Captive | Adult | Handling: mild/severe<br>(20 min) | Eye<br>Face<br>Wattle<br>Comb | ↓<br>=<br>↓<br>↓ | -0.4°C<br>=<br>-0.7 / -1.3°C<br>-0.5 / -2.2°C | (Herborn et al.,<br>2015) |
| Chicken<br>*Gallus gallus domesticus* | Captive | Adult | Enrichment removal<br>(within 2 h) | Eye<br>Face<br>Comb | ↓<br>↓<br>↓ | -0.8°C<br>-0.7°C<br>-3.5°C | (Herborn et al.,<br>2018) |
| Chicken<br>*Gallus gallus domesticus* | Captive | Chick<br>(30 days) | Handling<br>(10 min) | Head<br>Feet | ↑<br>↓ | +0.76°C<br>-0.45°C | (Moe et al., 2017) |
| Chicken<br>*Gallus gallus domesticus* | Captive | Chick<br>(9 days) | Visual<br>(45 min) | Eye<br>Beak | =<br>=, ↑ | =<br>+0.5°C | (Pijpers et al.,<br>2022) |
| Domestic pigeon<br>*Columbia livia domestica* | Captive | Adult | Handling<br>(3.5 min) | Eye<br>Beak | =<br>↓ | -0.4°C<br>-2.1°C | (Tabh et al., 2021) |
| House sparrow<br>*Passer domesticus* | Captive | Adult | Handling and injection<br>(~1min) | Eye<br>Beak | ↓<br>↓ | -1.0°C<br>-4.0°C | (Jerem and<br>Romero, 2023) |
| King penguin<br>*Aptenodytes patagonicus* | Wild | Adult | Handling<br>(5-10 min) | Eye<br>Beak | ↑<br>↓ | +0.8°C<br>-1.5°C | This study |
|  | Wild | Chick<br>(20 days*) | Handling<br>(5-10 min) | Eye<br>Beak | ↓<br>↓ | -0.9°C<br>-3.3°C |  |
| Little Auk<br>*Alle alle* | Wild | Adult | Handling (30 min) | Eye | ↑ | +1,7°C | (Jakubas et al.,<br>2022) |
| Pheasant<br>*Phasianus colchicus* | Captive | Juvenile<br>(42 to 49 days) | Agression<br>(20 s) | Head | ↑ | +0.6°C | (Knoch et al.,<br>2022) |
| Song sparrow<br>*Melospiza melodia* | Captive | Adult | Handling<br>(13 min) | Eye<br>Beak | ↓, =<br>↓, ↑, = | -1.5°C, =<br>-2.6°C, +2.6°C,<br>= | (Zuluaga and<br>Danner, 2023) |

\* king penguins chicks are not thermally emancipated at this age

In this study, we took up the challenge of distinguishing the role of HPA and SAM axes on changes in body surface temperatures using an experimental approach in adult king penguins (*Aptenodytes patagonicus*). Breeding king penguins were subcutaneously implanted with either a corticosterone (CORT, the main GC hormone in avian species) implant or a placebo implant. The use of CORT implants in king penguin (see methods and Fig. 1), as in many other bird species (see Torres-Medina et al., 2018), leads to higher baseline CORT levels, while inhibiting the acute CORT response in the days following implantation. Indeed, CORT-implanted birds no longer exhibit high blood CORT release in response to acute stress induced by a handling stress protocol because of the HPA negative feedback loop (Torres-Medina et al., 2018). Specifically, high circulating CORT levels are inhibiting the activity of the paraventricular nucleus in the hypothalamus through binding with glucocorticoid receptors. This inhibits the release of corticotrophin-releasing hormone and the afferent HPA cascade through the pituitary (secreting the adrenocorticotropic hormone, ACTH) and adrenal (secreting CORT) glands (Smulders, 2021). Hence, the use of CORT implants in conjunction with handling stress allows for the separation of the contributions of the HPA and SAM axes to acute changes in body surface temperatures. Here, we therefore measured the effects of our CORT treatment, together with a handling stress protocol, on surface temperatures in the king penguin. If changes in surface temperatures are mainly driven by activating the HPA axis, we expect to find higher initial body surface temperatures in CORT-implanted individuals (with elevated baseline CORT) prior to capture and handling. However, because CORT implants shut-down the release of CORT in response to handling, we expect to find no change in body surface temperature in response to handling stress in CORT individuals if such changes are mediated by the HPA axis only. On the contrary, if changes in surface temperatures are mainly driven by the activity of the SAM axis, we expect our CORT treatment to have no effect on body surface temperatures (both before or after capture and handling).

**Fig. 1:**
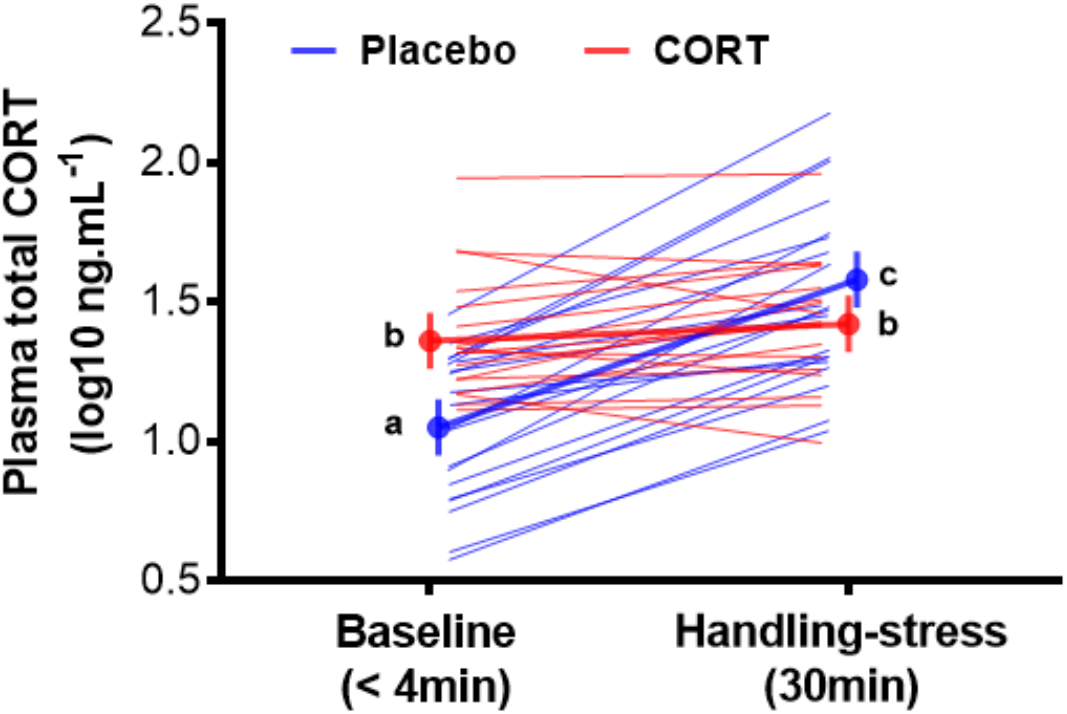
Baseline and stress-induced plasma corticosterone levels of breeding king penguins. implanted either with a Placebo, or a CORT subcutaneous implant (50 mg, 90 days release; n = 80 observations, N = 40 individuals) during incubation. Birds were blood sampled 6 days after implantation. CORT (red) *vs.* Placebo (blue) individual responses (raw data) are presented along with group and time specific mean ± SE. CORT levels were influenced by the interaction between handling stress (baseline: < 4 min vs. stress-induced: 30 min of handling stress) and CORT treatment (LMM on log-transformed plasma CORT: F_1,37.2_ = 62.4, *p* < 0.001). Letters indicate significant differences according to post-hoc tests with Tukey adjustments (*p* < 0.030).

In addition to direct effects on adults, parental stress exposure or increased CORT levels may also affect the following generation, for instance by affecting parental care (Thierry et al., 2013a) or by conveying information about a stressful environment (Brandl et al., 2022). The latter idea has been framed as the concept of family-transmitted stress by Noguera et al. (2017). Therefore, we also aimed to investigate whether thermal imaging could be used to quantify indirect effects of stress on king penguin’s progeny. To this end, we measured egg surface temperature on the same day as the body surface temperatures of implanted parents, and the surface temperatures of their chicks. Based on experimental data in Adélie penguin (*Pygoscelis adeliae*) showing that elevated baseline CORT levels in adults led to lower parental care (Thierry et al., 2013a), we hypothesized that increasing CORT levels may lower the surface temperatures of eggs and/or chicks during incubation and brooding respectively, as parents may pay less attention in keeping the egg/chick covered and warm under their brood pouch.

## 2. Material and Methods

### 2.1. Study site and species

The king penguin is a heterothermic bird whose body temperature largely varies between core and peripheral tissues (Lewden et al., 2017a; Lewden et al., 2017b). Its thick body plumage provides a very efficient insulation, while counter-current vascular heat exchangers in the appendages allows for a decrease in peripheral temperature loss (Thomas et al., 2011). These adaptations allow birds to conserve heat when foraging in cold Antarctic waters (Handrich et al., 1997), and to rapidly recover to normothermia when exiting the water (Lewden et al., 2020).

The study was conducted in 2018-2019 in a king penguin colony of *ca.* 22,000 breeding birds at ‘La Baie du Marin’ on Possession Island (Barbraud et al. 2020), Crozet Archipelago, in the Southern Indian Ocean (46°26’ S, 51°52’ E). We followed 49 breeding pairs from courtship (early November) until the onset of the Austral winter (April). All adults were identified by a hair dye mark on the breast feathers. During the early breeding season, male and female alternate between periods on-land caring for their single-egg or chick, and periods foraging at sea (Weimerskirch et al., 1992). The male takes the first incubation shift while the female forages at sea. The female returns ∼15 days later to relieve her partner, and the parents continue to alternate shifts throughout incubation (∼53 days) and early chick brooding. Chicks become thermally emancipated from the parents around one month of age, allowing both parents to go back at sea simultaneously to forage (Weimerskirch et al., 1992). Twenty days after hatching, king penguin chicks are able to maintain their core body temperature under mild environmental conditions (> 10°C) but still rely on their parent’s brooding behaviour (Duchamp et al., 2002), and are able to express a HPA-dependent stress response (X et al. unpublished).

### 2.2. Use of sub-cutaneous CORT and SHAM implants

We divided the 49 breeding pairs into two groups of either female-treated pairs (N = 23) or male-treated pairs (N = 26). In the female-treated pairs, we implanted females with either a CORT (NG-111, 50 mg corticosterone, 90-days release, 8 mm diameter and 3mm thickness; N = 12 females) or a Placebo (NC-111, same vehicle but no CORT; N = 11 females) implant. Sub-cutaneous implants were purchased from Innovative Research of America (Sarasota, FL, USA). Females were implanted during their first incubation shift, 3 days after returning from sea. Similarly, in the male-treated pairs, we implanted 13 males with a CORT and 13 males with a Placebo implant, 3 days after returning from sea. In males, the first incubation shift is immediately after egg laying, and males have already been fasting for ∼15 days during courtship. Therefore, we chose to implant the males during their second incubation shift so that they would have comparable fasting times compared to females, and therefore both sexes would be in a comparable physiological state during measurements. Allocation of birds to specific groups was designed to avoid spatial or temporal biases within and between groups. Male and female partners in, respectively, the female-treated and male-treated pairs were not implanted. Hereafter, they are referred to as “Partner-CORT” or “Partner-Placebo” regardless of their sex. Partner penguins were only considered in the chick analysis (see below).

On the day of the implantation, adults were captured by hand while incubating their egg and restrained by one experimenter with a hood covering the head to keep them calm. While maintaining individuals on their territory within the colony to avoid risking reproductive abandonment, sub-cutaneous CORT or placebo implants were inserted in the upper part of the back by a veterinary surgeon under local anaesthesia (*ca.* 0.5 mg/kg xylocaine combined with 0.0001 mg/kg adrenalin, Aspen Pharma). Immediately after implantation, the small incision (*ca.* 15 mm) was closed using 3 sterile surgical staples (SurgiClose^TM^ Skin Stapler), and a prophylactic dose of antibiotics (cephalexin *ca.* 50 mg/kg, Rilexine®, Virbac) was injected to prevent any risk of infection. The incision was checked 3 days later for implant rejection; no sign of infection or implant rejection were noticed and staples were removed 6 days after implantation.

To verify the efficacy of our CORT treatment, 6 days after implantation, 1 mL of blood was sampled from the flipper vein using a heparinized syringe in less than 4 min after being detected by the focal individual (baseline CORT) and after 30 min of standardized handling (stress-induced, see Stier et al. (2019) and Viblanc et al. (2018)). Blood samples were taken between 09:00 and 13:00 to limit variation linked to circadian variation in plasma CORT levels. Blood samples were centrifugated within one hour of sampling at 3000 *g* for 10 minutes. The plasma fraction was then removed and plasma aliquots were stored at −20°C until the end of the day, before being transferred to −80°C until laboratory analyses in 2023. Plasma total CORT levels were measured from 25 μL of plasma by immunoassay according to guidelines provided by the manufacturer (Corticosterone Enzyme Immunoassay Kit, Arbor Assay, USA). Intra-plate coefficient of variation based on duplicates was 8.17 ± 0.90% (mean ± SE), and inter-plate coefficient of variation based on one repeated sample was 8.08%. We tested the effect of implant on log-transformed plasma CORT using a linear mixed model (LMM) including as fixed factors: Handling stress (Baseline *vs.* Handling stress), Treatment (CORT *vs.* Placebo) and their interaction, as well as penguin ID as random effect. Implants were successful in raising circulating baseline CORT levels and in inhibiting stress-induced CORT release (Fig. 1).

### 2.3. Thermal image collection and analysis

Because of their dense and watertight plumage, the use of thermal imaging in seabirds is usually restricted to measuring the eye (periorbital region) and beak regions (Gauchet and Grémillet, 2022). These regions are highly vascularised and provide valuable information on body surface temperatures. Thermal pictures of implanted adults (Fig. 2A) and their egg (Fig. 2B) were taken 3 days after implantation and thermal pictures of chicks (Fig. 2C) were taken at day 20 after hatching (mean ± SE: 49.8 ± 1.4 days after CORT or Placebo implant of the focal parent, *i.e.* within the 90-days release given by the CORT pellet manufacturer). Thermal pictures were taken at a distance between *ca.* 0.8 and 2 m using FLIR E8 thermal imaging camera (320 x 240 pixels) resulting in a spotsize between 1.95 and 4.88 mm (Playa-Montmany and Tatersall 2021). King penguins’ beaks being approximately 120 mm long, and the eye region being approximately 18 mm of diameter, our target areas exceeded the 3 times larger spotsize recommended by Playa-Montmany and Tatersall (2021). When approaching the adults and/or chicks, we started a stopwatch when the focal individual showed the first signs of alarm behaviour (*e.g.* stopping activity, looking towards the experimenter) to measure the duration of our disturbance at the time the different pictures were taken. In adults, we took a first set of pictures (1 to 2 pictures per individual) before handling, on average (mean ± SE) 0.63 ± 0.07 min after the first sign of alarm behaviour at a distance of *ca*. 1.5 to 2 m from the focal individuals. Adults were then captured-restrained, incision healing was checked, and we took a second set of pictures (1 to 2 pictures per individual) after terminating handling stress (mean ± SE handling stress duration was 8.3 ± 0.3 min; a timeframe within which CORT is known to increase by *ca.* 100% above baseline levels in king penguins (Viblanc et al., 2018).

**Fig. 2:**
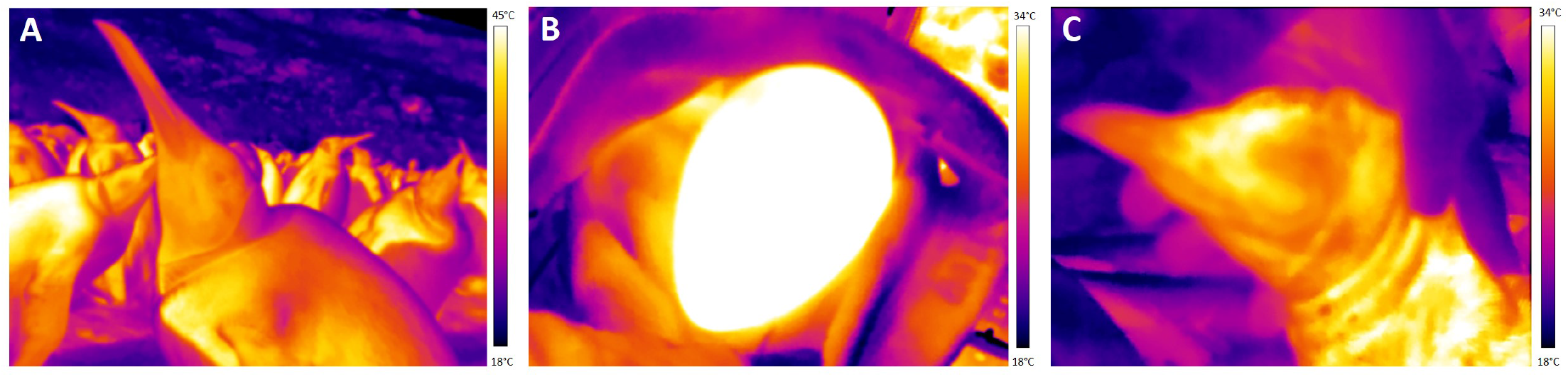
Infrared images of (A) breeding adult, (B) incubated egg and (C) 20 days-old chick.

Just after capturing the adult, we removed the single-egg from the brood pouch and immediately (*ca.* within 10 s) took a thermal picture at a distance of *ca.* 0.8 m. We then measured egg length and breadth using a Vernier calliper (Mitutoyo; accuracy ± 0.02 mm) and estimated egg volume (cm^3^) following (Narushin, 2005). When chicks were 20 days old, we took a first set of pictures (1 to 2 pictures per individual) at a distance of *ca.* 0.8 m immediately (*ca.* within 10 s) after they were removed from the brood pouch (*i.e.* before handling: 0.73 ± 0.08 min after the first sign of alarm behavior from the parent) and a second set of pictures (1-2 pictures per individual) was taken after handling, 5.40 ± 0.30 min later. Thermal images of the chicks were only taken on non-rainy days, which reduced the sample size from 38 chicks that survived until day 20, to 16 chicks with thermal images available. Chick sex was unknown and body mass was not significantly influenced by the parental treatment (CORT: 1.14 ± 0.11 kg *vs.* Placebo 1.32 ± 0.09 kg, N = 8/group, t-test: *t* = –1.28, *p* = 0.22). Chicks were kept within the colony at ambient temperature (14.9 ± 0.9°C) between the first and second set of pictures, a temperature at which they are known to be able to maintain their internal body temperature (Duchamp et al., 2002). In adults and chicks, we defined ‘before handling’ surface temperatures, the temperatures extracted from the first set of pictures taken in < 1 min and ‘after handling’ surface temperatures, the temperatures extracted from the second set of pictures taken after > 5 minutes of handling stress.

We analysed thermal images using the ThermaCAM TM Researcher Pro 2.10 software (Flir systems, Wilsonville, Oregon, USA). Only profile pictures were analysed to avoid surface temperature errors related to head orientation (Playà-Montmany and Tattersall, 2021; Tabh et al., 2021). For each image we set the emissivity at 0.98, reflected temperature at 20°C and controlled in the analyses for daily variation in air temperature (*T*_a_) and relative humidity using daily climatic measures from a permanent Météo France weather station located 2 km inland from the colony (https://rp5.ru/Archives_m%C3%A9t%C3%A9o_sur_la_base_Alfred-Faure). We extracted measures of surface temperatures on maximum periorbital temperature (*T*_eye_) and minimum beak temperature (*T*_beak_) in adults and chicks. We used maximum *T*_eye_ as recommended by Jerem et al. (2015 & 2019) as well as recorded in recent studies (*e.g.* Robertson et al. 2020a; Zuluaga & Danner 2023, and the minimum *T*_beak_ to gain insight on the maximum state of vasoconstriction for this body surface (Tattersall et al., 2009). Finally, we estimated egg surface temperature (*T*_egg_) as the mean of average length and width surface temperature.

### 2.4. Ethical note

All the procedures were approved by the French Ethical Committee (APAFIS#16465-2018080111195526 v4) and the Terres Australes et Antarctiques Françaises (Arrêté TAAF A-2018-118).

### 2.5. Statistical analysis

We ran separate analyses for adults, eggs and chicks. In adults, we investigated variations in *T*_eye_ and *T*_beak_ using two separated linear mixed models (LMMs) where Treatment (CORT *vs.* Placebo), Handling stress (before *vs.* after), Sex and *T*_a_ were specified as fixed effects. Relative humidity was initially tested but removed from final models since it was never significant. All two-ways interactions were also initially included but removed from the final model if *p* > 0.10 using a backward stepwise procedure. The p-values (just before removal in the backward stepwise procedure) for the focal interaction between Treatment and Handling stress are reported in tables. Bird identity was included as a random intercept to control for repeated measures. We investigated variation in *T*_egg_ by entering parental CORT Treatment (CORT *vs.* Placebo), the Sex of the parent, *T*_a_ and egg volume as fixed effects. Eggs were always measured when incubated by the implanted parents (*i.e.* 3 days after implantation of CORT or Placebo implants), while chicks at day 20 could be brooded by the implanted parent or its non-implanted partner. Thus, we investigated variation chick *T*_eye_ and *T*_beak_ by entering parental Treatment (CORT *vs.* Placebo), the category of the brooding parent at the time of measurement (implanted individual *vs.* non-implanted partner), Handling stress (before *vs.* after), and *T*_a_ as fixed effects, and chick identity as a random intercept. Parental sex was not included in the models for chick *T*_eye_ and *T*_beak_ due to limited sample size available for chicks. Interactions were treated as described above for adults. All statistical analyses were conducted using *R* (version 3.5.3) and the packages *lme4* (Bates et al., 2015), *emmeans* (Lenth et al., 2018) and *pbkrtest* (Halekoh and Højsgaard, 2014). Results are reported as least-square means ± SE.

## 3. Results

### 3.1. Adult surface temperatures

Treatment (CORT *vs.* Placebo), either alone or in interaction with Handling stress, had no significant effect on eye (*T*_eye_) or beak (*T*_beak_) surface temperature in incubating adults (Table 2, Fig. 3A and 3B). Handling stress significantly affected *T*_eye_ and *T*_beak_, in opposite directions (Table 2). Over an 8-min handling stress, *T*_eye_ increased by 0.8 ± 0.3°C, on average (Fig. 3A), while *T*_beak_ decreased by –1.5 ± 0.4°C, on average (Fig. 3B). The increase in *T*_eye_ appeared to be sex-specific (marginally significant interaction Handling * Sex: *p* = 0.060), with only females showing a significant increase in *T*_eye_ in response to Handling stress (Fig. 3C; males: *t* = 0.63, *p* = 0.53; females: *t* = 3.08, *p* = 0.003). As expected, *T*_eye_ and *T*_beak_ were significantly and positively associated with *T*_a_ (Table 2).

**Fig. 3:**
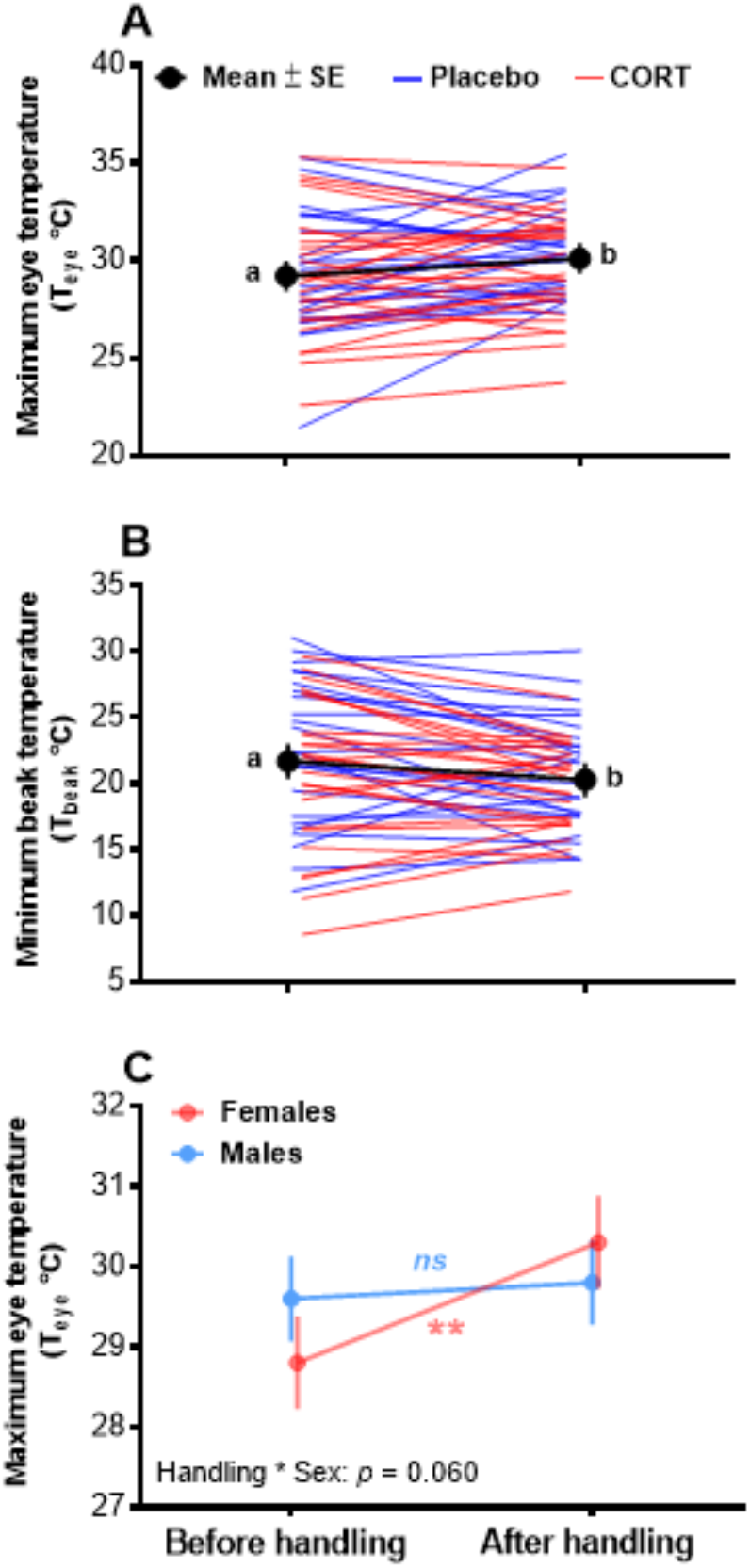
Adult king penguin surface temperature responses to an experimental manipulation of glucocorticoid levels (CORT *vs.* Placebo subcutaneous implants) and handling stress: (A) Maximum eye temperature response to handling stress, (B) Minimum beak temperature response to handling stress, and (C) Sex-specific response of eye temperature to handling stress. In panels A and B, CORT (red) *vs.* Placebo (blue) individual responses (raw data) are presented. Since CORT <u>T</u>reatment had no significant effect either alone or in interaction with Handling stress (see Table 2 for details), the overall mean ± SE is shown in black. Least-square means ± SE from final statistical models (Table 2; n = 138 observations; N = 49 individuals) are presented. Different letters indicate significant differences in panels A and B, and ** in panel C represent the significant increase of T_eye_ observed only in females.

**Table 2:** Summary of linear mixed models (LMMs) investigating the determinants of (A) maximum eye temperature (*T*_eye_) and (B) minimum beak temperature (*T*_beak_) in adult king penguins.

**A. $T_{eye}$ (n = 138 observations; N = 49 individuals)**
| Random effects: |  | Variance |  |  |  |
| --- | --- | --- | --- | --- | --- |
| Bird ID | Intercept | 5.04 |  |  |  |
| Residual |  | 3.21 |  |  |  |
| Fixed effects: | Estimate | SE | t | df | p |
| Intercept | 24.94 | 1.71 | 14.54 | 47.9 | <0.001 |
| Treatment (CORT) | -0.10 | 0.72 | -0.15 | 45.8 | 0.89 |
| **Handling (after)** | **1.45** | **0.47** | **3.04** | **89.3** | **0.003** |
| Sex (Male) | 0.76 | 0.78 | 0.97 | 63.7 | 0.34 |
| $T_a$ | **0.29** | **0.13** | **2.33** | **45.0** | **0.024** |
| *Handling\*Sex* | -1.19 | 0.63 | -1.90 | 89.7 | 0.060 |
| (Treatment x Handling) |  |  |  |  | (0.44) |

**B. $T_{beak}$ (n = 138 observations; N = 49 individuals)**
| Random effects: |  | Variance |  |  |  |
| --- | --- | --- | --- | --- | --- |
| Bird ID | Intercept | 15.91 |  |  |  |
| Residual |  | 6.40 |  |  |  |
| Fixed effects: | Estimate | SE | t | df | p |
| Intercept | 13.84 | 2.92 | 4.74 | 45.0 | <0.001 |
| Treatment (CORT) | -1.13 | 1.24 | -0.91 | 44.5 | 0.37 |
| **Handling (after)** | **-1.47** | **0.45** | **-3.29** | **84.6** | **0.001** |
| Sex (Male) | 0.90 | 1.23 | 0.73 | 44.6 | 0.47 |
| $T_a$ | **0.54** | **0.22** | **2.45** | **43.8** | **0.018** |
| (Treatment x Handling) |  |  |  |  | (0.61) |

### 3.2. Egg surface temperature

Neither parental Treatment (*F*_1,45_ = 1.38, *p* = 0.25; CORT: 34.5 ± 0.2°C *vs.* Placebo: 34.9 ± 0.3°C; N = 46), Sex (*F*_1,45_ = 0.47, *p* = 0.49), ambient temperature (*F*_1,45_ = 0.45, *p* = 0.50), nor egg volume (*F*_1,45_ = 0.01, *p* = 0.93) significantly affected *T*_egg_.

### 3.3. Chick surface temperatures

The interaction between parental Treatment (CORT *vs.* Placebo) and the category of the parent brooding the chick (implanted *vs.* non-implanted partner) at the time of measurement significantly affected chick *T*_eye_ (*F*_1,11_ = 6.37, *p* = 0.028, Table 3) and chick *T*_beak_ (*F*_1,11_ = 5.49, *p* = 0.039, Table 3). Chicks brooded by the partner of CORT-implanted individuals had higher overall *T*_eye_ and *T*_beak_ than chicks brooded by CORT parents, or by Placebo and associated partners (Fig 4A and 4B). While we found no significant interaction between parental Treatment and Handling stress (Table 3), both chick *T*_eye_ (Fig 4C) and chick *T*_beak_ (Fig 4D) significantly decreased after handling (by –0.9 ± 0.3°C and –3.3 ± 0.6°C respectively, Table 3). As expected, chick *T*_eye_ and *T*_beak_ were significantly positively associated with T_a_ (Table 3).

**Fig. 4:**
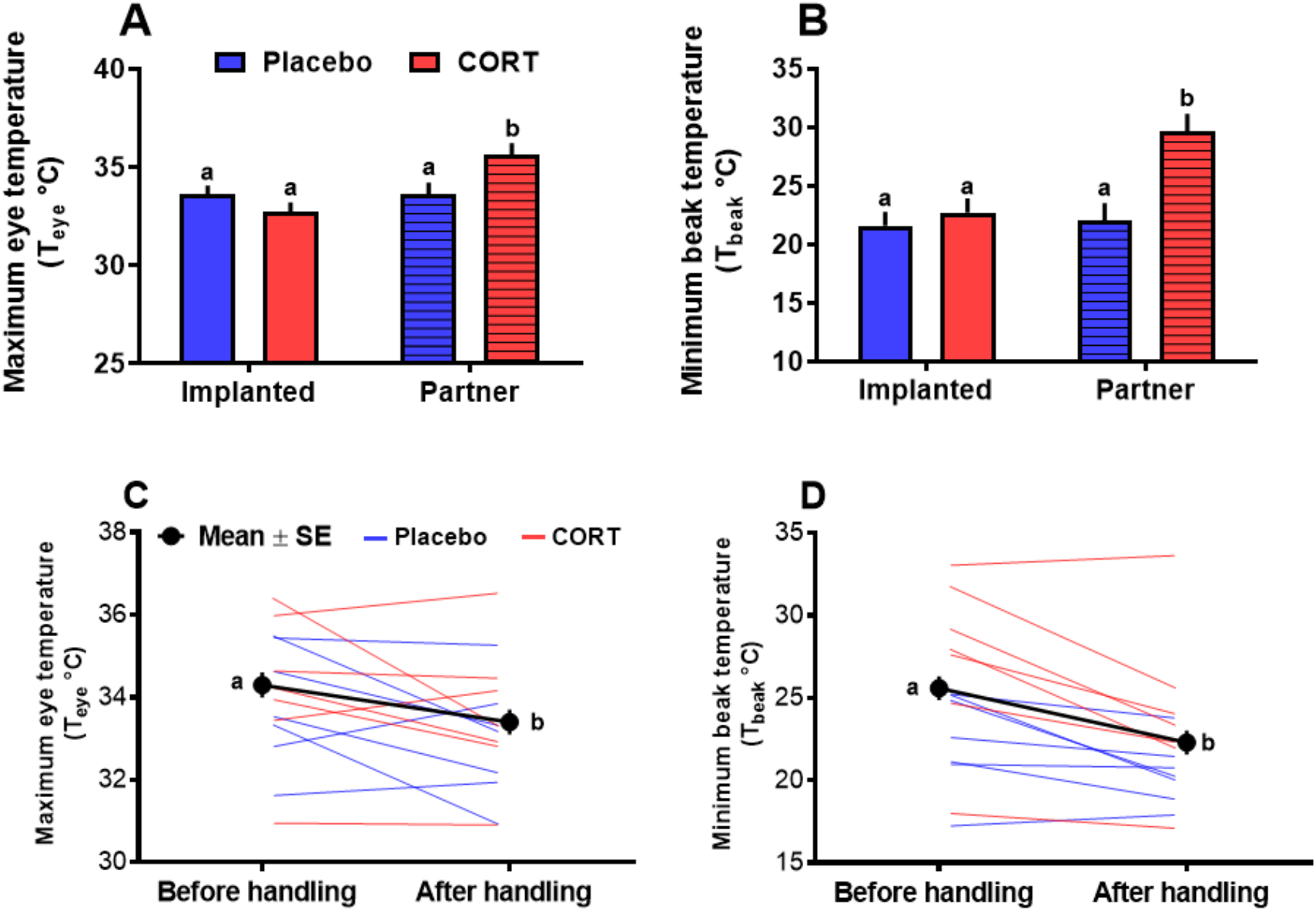
King penguin chick surface temperature in response to parental glucocorticoid manipulation and handling: (A) Maximum eye temperature according to parental treatment and category of the parent brooding the chick (*i.e.* partners are not implanted), (B) Minimum beak temperature according to parental treatment and category of the parent brooding the chick, (C) Maximum eye temperature response to handling, and (D) Minimum eye temperature response to handling. In panels C and D, CORT (red) vs. Placebo (blue) chick’s individual responses (raw data) are presented, but only the global mean ± SE are presented since there was no significant interaction between parental CORT Treatment and Handling (see Table 3 for details). Least-square means ± SE from final statistical models (Table 3) are presented (n = 63 observations, N = 16 chicks). Different letters indicate significant differences.

**Table 3:** Summary of linear mixed models (LMMs) investigating the determinants of. (A) maximum eye temperature (*T*_eye_) and (B) minimum beak temperature (*T*_beak_) in king penguin chicks. Only one parent was implanted with CORT or Placebo implant, thus we considered both parental treatment (CORT *vs.* Placebo), the category of the parent brooding the chick at the time of measurement (implanted parent *vs.* non-implanted partner) and their interaction in our analyses.

**A. $T_{eye}$ (n = 63 observations; N = 16 chicks)**
| Random effects: |  | Variance |  |  |  |
| --- | --- | --- | --- | --- | --- |
| Bird ID | Intercept | 0.81 |  |  |  |
| Residual |  | 1.18 |  |  |  |
| Fixed effects: | Estimate | SE | t | df | p |
| Intercept | 29.47 | 1.39 | 21.26 | 11.1 | <0.001 |
| Treatment (CORT) | -0.88 | 0.68 | -1.29 | 11.3 | 0.22 |
| Parent (implanted) | **-2.84** | **0.82** | **3.48** | **11.1** | **0.005** |
| Handling (after) | **-0.85** | **0.28** | **-3.08** | **47.2** | **0.003** |
| $T_a$ | **0.21** | **0.09** | **2.41** | **10.9** | **0.035** |
| Treatment*Parent | **2.78** | **1.10** | **2.53** | **11.0** | **0.028** |
| (Treatment x Handling) |  |  |  |  | (0.79) |

**B. $T_{beak}$ (n = 63 observations; N = 16 chicks)**
| Random effects: |  | Variance |  |  |  |
| --- | --- | --- | --- | --- | --- |
| Bird ID | Intercept | 5.53 |  |  |  |
| Residual |  | 4.89 |  |  |  |
| Fixed effects: | Estimate | SE | t | df | p |
| Intercept | 13.55 | 3.42 | 3.96 | 11.1 | 0.002 |
| Treatment (CORT) | 1.15 | 1.68 | 0.69 | 11.3 | 0.51 |
| Parent (implanted) | **-6.82** | **2.02** | **-3.38** | **11.1** | **0.006** |
| Handling (after) | **-3.26** | **0.56** | **-5.77** | **46.9** | **<0.001** |
| $T_a$ | **0.57** | **0.21** | **2.65** | **11.0** | **0.023** |
| Treatment*Parent | **6.37** | **2.72** | **2.34** | **11.0** | **0.039** |
| (Treatment x Handling) |  |  |  |  | (0.94) |

## 4. Discussion

By experimentally manipulating CORT levels in adult breeding king penguins, we show that corticosterone is unlikely to play a major role in determining changes in surface temperatures (both before or in response to handling stress) in king penguins. Indeed, we found no significant difference in baseline surface temperature between CORT and Placebo implanted individuals. However, whatever the parental CORT treatment, our handling stress protocol led to an increase in *T*_eye_ and a decrease in *T*_beak_ in response to capture and handling. The absence of effect of the CORT treatment on baseline surface temperature and its dynamics during acute handling stress suggest that changes in surface temperature are probably driven primarily by the activity of the SAM axis in king penguins (but see below for a potential causal involvement of CORT in house sparrow *Passer domesticus* (Ouyang et al., 2021)). Contrary to our prediction, parental treatment had no significant effect on egg temperature. Yet, we found an unexpected pattern for chick surface temperatures: chicks brooded by the non-implanted partner of CORT individuals had higher surface temperatures (both *T*_eye_ and *T*_beak_) than chicks brooded by Placebo (or associated non-implanted partner) or CORT individuals. Finally, in the thermally non-emancipated chicks, both *T*_eye_ and *T*_beak_ decreased with handling, but irrespectively of the parental treatment.

### 4.1. CORT and adult surface temperatures

Previous correlative studies have considered baseline and stress-induced CORT levels to investigate the relationship between GC signalling and changes in body surface temperature in birds (blue tit, *Cyanistes caerulescens*, (Jerem et al., 2018); domestic chicken, *Gallus gallus domesticus,* (Giloh et al., 2012; Weimer et al., 2020)). To go deeper in the involved mechanisms, we experimentally manipulated CORT in the king penguin to help to causally distinguish the contribution of the HPA and SAM axes to changes in body surface temperature. Since we did not detect any significant effect of CORT manipulation on baseline or stress-induced surface temperatures, our results provide little support for a causal involvement of the HPA axis in mediating stress-related changes in surface temperature.

Our results contrast with a recent study showing that an acute stimulation of the HPA axis with adrenocorticotropic hormone (ACTH) induced a decrease in *T*_eye_ within 5 min in captive-held house sparrows (Ouyang et al., 2021), which suggests a causal involvement of the HPA axis in mediating stress-related changes in surface temperature. Yet, a recent study in the same species found a significant correlation between the acute change in surface temperature and heart rate variability, a measure of SAM axis activation (Jerem and Romero, 2023). This suggests that both the HPA and SAM axes could be involved in mediating stress-related changes in surface temperature in house sparrow. While we used an experimental approach and the lack of significant effect of baseline CORT on body surface temperatures appears to be robust, we cannot fully exclude that our handling stress protocol for thermal imaging was too short to detect a potential role of stress-induced CORT in influencing changes in surface temperatures. Indeed, in 8 minutes of handling, CORT levels are expected to double in king penguin, while maximum CORT levels are only attained after *ca.* 80 minutes of handling in our study species (Viblanc et al., 2018). Additionally, we may have missed some early changes in surface temperatures occurring in response to the stress of our approach prior to bird capture and handling. Indeed, changes in *T*_eye_ and *T*_beak_ within 25 seconds before capture have been observed in house sparrow (*Passer domesticus*) (Jerem and Romero, 2023) and our pre-handling thermal pictures were taken on average 38 seconds after the first sign of alarm behaviour. Yet, our results suggest that changes in surface temperature are probably more likely mediated by the SAM axis rather than the HPA axis in adult king penguins. Future studies testing for a direct acute stimulation of the HPA axis (using ACTH) or SAM axis (using catecholamines) in the king penguin and other bird species could be useful to gain greater insight on the contribution of the HPA and SAM axes in mediating changes in body surface temperature (Jerem and Romero, 2023; Ouyang et al., 2021).

### 4.2. Stress-induced changes in surface temperatures

In breeding adults, we observed a mild sex-specific significant increase of *T*_eye_ and a significant decrease of *T*_beak_ in response to acute capture-handling stress. Differences in temperature change between body regions in response to acute stress are not rare (see Table 1). For instance, (Moe et al., 2017) measured a decrease of *T*_eye_ contrasting with an increase of *T*_feet_ in chicken, whereas in captive budgerigars (*Melopsittacus undulatus*) *T*_eye_ increased transiently after exposure to a stressor while *T*_feet_ did not change (Ikkatai and Watanabe, 2015; Table 1). Anatomical and functional reasons can likely explain the difference of stress-induced changes in surface temperature between body regions in penguins, with for instance a potential interest in maintaining or increasing eye blood flow to maintain or enhance visual acuity when acutely threatened by an environmental challenge.

*T*_eye_ has also been shown to positively correlate with internal (cloacal) temperature in various avian species (budgerigars: Ikkatai and Watanabe, 2015; chicken: Cândido et al., 2020). Consequently, it is possible that the increase in *T*_eye_ we observe in adults could reflect an increase in internal body temperature (Oka, 2018). Consistently, in closely related Emperor penguins (*Aptenodytes forsteri*), acute stress leads to an approximate 1.5°C increase in internal (stomach) temperature (Regel and Pütz, 1997). Such an increase in *T*_eye_ in response to handling stress has been reported two times in avian species to the best of our knowledge (Ikkatai and Watanabe, 2015; Jakubas et al., 2022), while most previous studies have shown either a significant decrease (4 studies) or no significant change (4 studies; Table 1). Those contrasted findings might be explained by at least three factors: 1. variation in body size, since smaller individuals/species are expected to favour internal heat conservation by reducing more markedly peripheral blood flow and surface temperature compared to large ones (*thermoprotective hypothesis*; Robertson et al., 2020a); 2. differences in the thermal environment, since it has been shown that during acute stress response, heat conservation is favoured below the thermoneutral zone while heat dissipation is favoured above its upper limit (Robertson et al., 2020a); 3. the various delays at which *T*_eye_ was measured in response to acute stress, since for instance the increase in *T*_eye_ found by Ikkatai and Watanabe (2015) was short-lived (visible 5 min after the stressor, but not later on during the 30 min measurement period). It is important to note that *T*eye and *T*_beak_ can show very rapid and non-linear changes in response to acute stress, that are likely to be missed by most studies not using continuous thermal image recording (Jerem et al., 2019; Jerem and Romero, 2023; Zuluaga and Danner, 2023). For instance, Zuluaga et al. (2023) found an initial drop in *T_beak_* following exposure to stressor, with a recovery to baseline levels after only 2 minutes, while Jerem et al. 2019 found an initial drop in *T_eye_* lasting less than 15s, followed by an increase above baseline from 15 to 75 seconds.

Although the interaction between handling stress and sex was only marginally significant (*p* = 0.060), it appeared that only females exhibited a noticeable increase in *T*_eye_ in response to handling stress in our study. Differences in body size or mass between sexes are very modest (*ca.* 5%) in king penguin (Kriesell et al., 2018), which thus seems unlikely to fully explain the sex-difference observed here. There are known sex-differences in stress physiology (Handa and McGivern, 2017), but sex differences in surface temperature changes induced by acute stress have been rarely investigated (but see Robertson et al. (2020a) for a result opposite to ours between sexes). We previously observed no sex effect in HPA axis responsiveness between males and females king penguins (Viblanc et al., 2016), but as mentioned above, the SAM axis is the likely driver of acute changes in surface temperatures. Consequently, this suggests that sexes might differ in the stress-sensitivity of their SAM axis, although this would need to be confirmed by direct measurements of SAM axis activity.

The decrease we observe in *T*_beak_ in response to acute stress in adults appears like the typical peripheral vasoconstriction response previously reported for instance by Tabh et al. (2021). Such peripheral vasoconstriction enables the redistribution of the peripheral blood circulation to internal organs and tissues, such as the brain or muscles, favouring their oxygenation and nutrition to sustain the fight-or-flight response. King penguins breed in a highly dense and aggressive colonial environment (up to 500 aggressive interactions per hour; (CôTé, 2000). Aggressive interactions, frequently leading to injuries, are known to result in increased heart rates (Viblanc et al., 2012). A decrease of *T*_beak_ during acute stress response likely reflects peripheral vasoconstriction, that is likely widespread to other peripheral body parts, which according to the *haemoprotective hypothesis* could help to reduce blood loss in case of injury (Robertson et al., 2020a).

The changes observed in surface temperatures in chicks differ from those found in adults. Indeed, we observed a significant decrease in chick’s *T*_eye_, whereas *T*_eye_ increased in adult females and stayed stable in adult males. While *T*_beak_ significantly decreased as observed in adults, the decrease was more pronounced in chicks (‒3.3 ± 0.6°C *vs.* –1.5 ± 0.4°C, in chicks and adults, respectively). This is likely explained by greater peripheral thermal losses in chicks suddenly exposed to an ambient temperature drop (*ca.* 35°C under the brood pouch of the adults *vs.* 15°C when taken out for measurement and handling), despite their ability to maintain their internal temperature at this stage at ambient temperature (Duchamp et al., 2002). The more pronounced drop in surface temperature we observed in chicks during acute handling could reflect their greater need to conserve heat at a relatively mild *T*_a_ compared to adults.

### 4.3. Impact of parental CORT on egg and chick surface temperature

Contrary to our prediction, we observed no significant impact of parental CORT treatment on egg temperature, suggesting little alteration of incubation behaviour by increased CORT levels, contrary to a previous study in Adelie penguin reporting a reduction of 1.3 ± 0.2°C in *T*_egg_ (Thierry et al., 2013a). It is possible that incubating two eggs in the colder environment of Adélie penguins is more challenging (and thus more likely to be impacted by high CORT levels) than incubating a single egg under the milder climate experienced by king penguins. Alternatively, it is also possible that king penguins are more resilient to increased CORT levels than Adelie penguins, which is supported by the good reproductive success of king penguins implanted with CORT (X et *al.* unpublished), contrary to what has been observed in Adelie penguins (Thierry et al., 2013b). Measurements of incubation quality using a dummy egg with temperature and rotation sensors (Thierry et al., 2013a) may provide more accurate data on this question, but it is important to note that the *T*_egg_ measured in this study (Placebo: 34.9 ± 0.3 °C) was close to the *T*_egg_ measured in the same penguin colony using internal sensors in dummy eggs (35.7 ± 0.4°C; (Groscolas et al., 2000)). Our results therefore suggest that further studies may benefit from the use of minimally invasive thermal imaging to measure incubation quality, for instance in the context of parental behaviour and climate change (Cook et al., 2020).

The measure of chick surface temperatures revealed some surprising patterns. While we expected CORT chicks to have lower surface temperatures due to poorer parental care (less efficient brooding), we found some opposite result: chicks had higher surface temperatures when brooded by the non-implanted partner of the CORT parent. This result should be considered with caution considering the limited sample size available for chick surface temperatures (N = 16). One possibility is that partners from CORT-implanted individuals somehow perceived the ‘stress’ levels of their partner (as shown between siblings in yellow legged gulls (*Larus michahellis*); Noguera et al., 2017) and somehow compensated for parental care through more efficient brooding. This hypothesis requires further study to determine whether the concept of ‘family-transmitted stress’ (Noguera et al., 2017) applies to the king penguin. In this species, both parents must rely on each other over more than one year to successfully fledge their single chick, which itself is entirely dependent on its parents for its food supply throughout its growth. Hence, the king penguin could be promising system to investigate the ‘family-transmitted stress’ hypothesis.

### 4.4. Conclusion

Evaluating stress levels and responses of free-living animals in a non-invasive manner using thermal imaging is a burgeoning field of research (Jerem et al., 2015; Tabh et al., 2022). However, little is still known about the underlying pathways of stress physiology that influence the changes in surface temperature. Our experimental study in king penguin points toward a likely preponderant role of the SAM axis. Additionally, by summarizing the current evidence for acute changes in surface temperatures during stress exposure (Table 1), we highlight the relative complexity and inconsistency between the effects observed by different studies. Further experimental studies related to SAM and HPA axes involvement are thus required to clarify what stress component(s) are measured through non-invasive thermal imaging.

## 5. Declarations

### 5.1. Author contribution

Study design: AS, VAV, PB and AL. Funding acquisition: JPR, PB, VAV, AS. Data collection in the field: AS, SA, TH, LA, CG, and JPR. Data collection in the lab: ER and AS. Data analysis: CW, AL, PB, AS. Writing original draft: AL, PB and AS. Writing review and editing: VAV, JPR, AN, SA, TH, LA, ER and CG.

### 5.2 Data availability

The datasets used in this manuscript are available on *FigShare* (10.6084/m9.figshare.20134181).

## Abbreviations

ACTH: Adrenocorticotropic hormone
*T*_a_: Ambient air temperature
CORT: Corticosterone
GC: Glucocorticoid hormones
*T*_egg_: Egg surface temperature
HPA: Hypothalamic-pituitary-adrenal
LMMs: Linear mixed models
*T*_eye_: Maximum eye region surface temperature
*T*_beak_: Minimum beak surface temperature
SAM: Sympathetic-adrenal-medullary

## 5.3. Acknowledgments

We are grateful to the French Polar Institute (IPEV) and the Terres Australes et Antarctiques Françaises for providing financial and logistical support for this study through the polar program #119 (ECONERGY). This study is part of the long-term Studies in Ecology and Evolution (SEE-Life) program of the CNRS. We wish to thank the Zone Atelier Antarctique et Terres Australes (ZATA) from the CNRS for financial support, and the members of the Alfred Faure field station for their help and support in the field. AL was supported by ISblue project, Interdisciplinary graduate school for the blue planet (ANR-17-EURE-0015) and co-funded by a grant from the French government under the program “Investissements d’Avenir” embedded in France 2030. AS was supported by a ‘Turku Collegium for Science and Medicine’ Fellowship, a Marie Sklodowska-Curie Postdoctoral Fellowships (#894963) and the IdEx Université de Strasbourg (*HotPenguin*). SA and CG were funded by the French Polar Institute.

